# Efficient incorporation and template-dependent polymerase inhibition are major determinants for the broad-spectrum antiviral activity of remdesivir

**DOI:** 10.1101/2021.10.14.464416

**Authors:** Calvin J. Gordon, Hery W. Lee, Egor P. Tchesnokov, Jason K. Perry, Joy Y. Feng, John P. Bilello, Danielle P. Porter, Matthias Götte

**Affiliations:** Department of Medical Microbiology and Immunology, University of Alberta, Edmonton, Alberta, Canada; Gilead Sciences, Inc., Foster City, California, USA

**Keywords:** Remdesivir, RNA virus, RNA-dependent RNA polymerase, broad-spectrum antiviral, COVID-19, SARS-CoV-2, Ebola, Nipah, Lassa, influenza

## Abstract

Remdesivir (RDV) is a direct antiviral agent that is approved in several countries for the treatment of coronavirus disease 2019 (COVID-19) caused by the severe acute respiratory syndrome coronavirus 2 (SARS-CoV-2). RDV exhibits broad-spectrum antiviral activity against positive-sense RNA viruses, e.g., SARS-CoV-2 and hepatitis C virus (HCV) and non-segmented negative-sense RNA viruses, e.g., Nipah virus (NiV), while several segmented negative-sense RNA viruses such as influenza (Flu) virus or Crimean-Congo hemorrhagic fever virus (CCHFV) are not sensitive to the drug. The reasons for this apparent pattern are unknown. Here, we expressed and purified representative RNA-dependent RNA polymerases (RdRp) and studied three biochemical parameters that have been associated with the inhibitory effects of RDV-triphosphate (TP): (i) selective incorporation of the nucleotide substrate RDV-TP, (ii) the effect of the incorporated RDV-monophosphate (MP) on primer extension, and (iii) the effect of RDV-MP in the template during incorporation of the complementary UTP. The results of this study revealed a strong correlation between antiviral effects and efficient incorporation of RDV-TP. Delayed chain-termination is heterogeneous and usually inefficient at higher NTP concentrations. In contrast, template-dependent inhibition of UTP incorporation opposite the embedded RDV-MP is seen with all polymerases. Molecular modeling suggests a steric conflict between the 1’-cyano group of RDV-MP and conserved residues of RdRp motif F. We conclude that future efforts in the development of nucleotide analogues with a broader spectrum of antiviral activities should focus on improving rates of incorporation while capitalizing on the inhibitory effects of a bulky 1’-modification.

## Introduction

The coronavirus disease 2019 (COVID-19) caused by severe acute respiratory syndrome coronavirus 2 (SARS-CoV-2) revealed the importance of broad-spectrum antivirals as the first line of defense in a pandemic(1,2). Vaccines that effectively protect against severe COVID-19 were developed in less than a year after the World Health Organization (WHO) declared a pandemic in March 2020(3–6). Antibody therapies, directed specifically against SARS-CoV-2, were also developed in a relatively short time and the first antiviral drug approved for the treatment of COVID-19 was the nucleotide analogue prodrug remdesivir (RDV)(7). Its broad spectrum of antiviral activity, including CoV’s, was described in earlier pre-clinical studies(8–17). RDV is a 1’-cyano modified *C*-adenosine monophosphate prodrug that targets the RNA-dependent RNA polymerase (RdRp) of a diverse panel of RNA viruses(8,13,18–20). In 2019, prior to the COVID-19 pandemic, RDV was tested in a randomized clinical phase 3 trial for the treatment of Ebola virus disease (EVD)(21). Although RDV’s clinical efficacy against EVD is inferior to two antibody therapies, its human safety data became available and this enabled RDV compassionate use in January 2020 to treat COVID-19 patients(22).

Clinical studies of RDV for the treatment of COVID-19 have shown a shortened time to recovery with an unclear effect on mortality (23,24). RDV is intravenously administered and is therefore limited to patients under supervised medical care. In general, oral antiviral drugs would allow earlier initiation of treatment in a broader patient population which may reduce the risk of progression to more severe disease outcomes, not only in the context of infection with coronaviruses, but potentially for the treatment of other viral infections. The development of oral broad-spectrum antiviral drugs is therefore critical to public health and pandemic preparedness. Successful efforts in this field require a better understanding of biochemical mechanisms that translate to antiviral activity. RDV shows antiviral activity against human positive-sense RNA viruses including members of *Coronoviridae* (SARS-CoV, Middle Eastern respiratory syndrome, MERS-CoV, and SARS-CoV-2)(8,19) and the *Flaviviridae* (HCV)(20), as well as non-segmented negative-sense RNA viruses including members of the *Filoviridae* (Zaire ebolavirus [Ebola virus, EBOV]), *Pneumoviridae* (respiratory syncytial virus [RSV]), and *Paramyxoviridae* (Nipah virus (NiV])(13). However, antiviral activity against segmented negative RNA viruses is less pronounced or not significant, as shown with members of the *Arenaviridae* (Lassa virus [LASV]), the *Orthomyxoviridae* (influenza [Flu] virus), and *Nairoviridae* (Crimean-Congo hemorrhagic fever virus [CCHFV])(13,20).

Here we expressed and purified representative RdRp enzymes or complexes associated with these viruses and studied biochemical mechanisms of RDV-mediated inhibition of RNA synthesis to better understand the molecular requirements for antiviral effects. Previous studies with purified SARS-CoV, MERS-CoV and SARS-CoV-2 RdRp complexes have shown that the triphosphate form of RDV (RDV-TP) is 2-3 times more efficiently incorporated than its natural counterpart ATP (25–28). Additionally, the incorporated monophosphate (RDV-MP) at position “i” causes delayed chain-termination at position “i+3”, which provides a possible mechanism for inhibition of RNA synthesis although increases in NTP concentrations can overcome this obstacle. However, under these circumstances, RDV-MP becomes embedded in the newly synthesized RNA that later serves as a template. Reduced rates of UTP incorporation opposite the complementary RDV-MP were shown to provide a second opportunity for inhibition(29). Our results demonstrate that the template-dependent inhibition of RNA synthesis is observed across a diverse panel of viral polymerases, which represents a necessary but not sufficient mechanism of action. Efficient, selective incorporation of RDV-TP is an important prerequisite for downstream inhibition of RNA synthesis and its translation into antiviral activity.

## Results

### Selective incorporation of RDV-TP

To address the differences seen in the antiviral activity of RDV against a spectrum of virus families, we compared the efficiency and pattern of inhibition of recombinant RdRp enzymes or enzyme complexes representing members of relevant families of viruses, and for which antiviral activity data were available (Table 1). Partial biochemical data were available for SARS-CoV, MERS-CoV, SARS-CoV-2, EBOV, and RSV RdRp complexes. Here we included enzymes and enzyme complexes of HCV and NiV that are also sensitive to RDV, LASV that is less sensitive to RDV, and influenza B (FluB) and CCHFV that are not sensitive to RDV by antiviral assay. RNA synthesis was monitored with short primer/templates mimicking random elongation complexes as previously described(16,25,26,29,30). We first determined the steady-state kinetic parameters for RDV-TP incorporation for each RdRp and normalized them to the rates of incorporation of ATP. The ratio of efficiency of single incorporations of ATP over RDV-TP defines the selectivity, which is a unitless parameter that facilitates comparisons between enzymes. Selectivity is less than 1, if incorporation of the analogue is more efficient than incorporation of its natural counterpart.

**Table 1.**
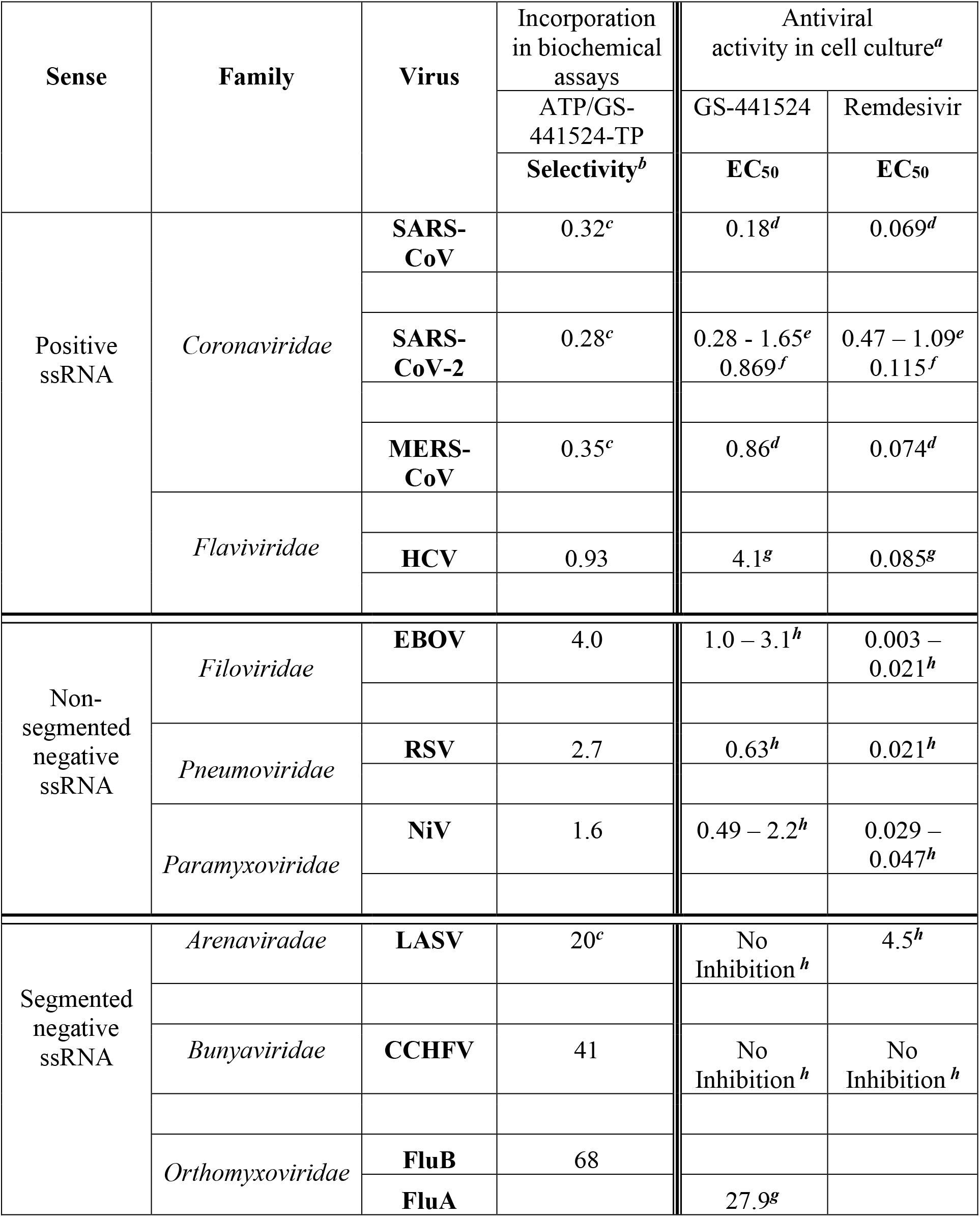
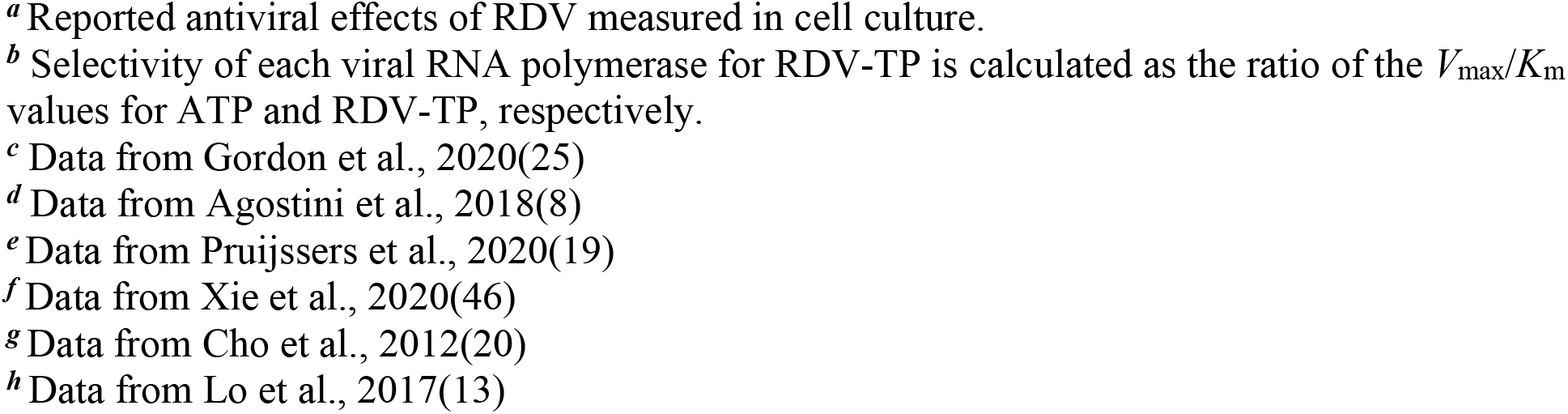
Selective incorporation of RDV-TP and antiviral activity against selected RNA viruses.

| Sense | Family | Virus | Incorporation in biochemical assays | Antiviral activity in cell culture<sup>a</sup> |  |
| --- | --- | --- | --- | --- | --- |
|  |  |  | ATP/GS-441524-TP | GS-441524 | Remdesivir |
|  |  |  | Selectivity<sup>b</sup> | EC<sub>50</sub> | EC<sub>50</sub> |
| Positive ssRNA | *Coronaviridae* | **SARS-CoV** | 0.32<sup>c</sup> | 0.18<sup>d</sup> | 0.069<sup>d</sup> |
|  |  | **SARS-CoV-2** | 0.28<sup>c</sup> | 0.28 - 1.65<sup>e</sup><br>0.869<sup>f</sup> | 0.47 – 1.09<sup>e</sup><br>0.115<sup>f</sup> |
|  |  | **MERS-CoV** | 0.35<sup>c</sup> | 0.86<sup>d</sup> | 0.074<sup>d</sup> |
|  | *Flaviviridae* |  |  |  |  |
|  |  | **HCV** | 0.93 | 4.1<sup>g</sup> | 0.085<sup>g</sup> |
| Non-segmented negative ssRNA | *Filoviridae* | **EBOV** | 4.0 | 1.0 – 3.1<sup>h</sup> | 0.003 – 0.021<sup>h</sup> |
|  | *Pneumoviridae* | **RSV** | 2.7 | 0.63<sup>h</sup> | 0.021<sup>h</sup> |
|  | *Paramyxoviridae* | **NiV** | 1.6 | 0.49 – 2.2<sup>h</sup> | 0.029 – 0.047<sup>h</sup> |
| Segmented negative ssRNA | *Arenaviridae* | **LASV** | 20<sup>c</sup> | No Inhibition<sup>h</sup> | 4.5<sup>h</sup> |
|  | *Bunyaviridae* | **CCHFV** | 41 | No Inhibition<sup>h</sup> | No Inhibition<sup>h</sup> |
|  | *Orthomyxoviridae* | **FluB** | 68 |  |  |
|  |  | **FluA** |  | 27.9<sup>g</sup> |  |

<sup>a</sup> Reported antiviral effects of RDV measured in cell culture.
<sup>b</sup> Selectivity of each viral RNA polymerase for RDV-TP is calculated as the ratio of the $V_{\max}/K_m$ values for ATP and RDV-TP, respectively.
<sup>c</sup> Data from Gordon et al., 2020(25)
<sup>d</sup> Data from Agostini et al., 2018(8)
<sup>e</sup> Data from Pruijssers et al., 2020(19)
<sup>f</sup> Data from Xie et al., 2020(46)
<sup>g</sup> Data from Cho et al., 2012(20)
<sup>h</sup> Data from Lo et al., 2017(13)

Previous data have shown that RdRp complexes of coronaviruses SARS-CoV, MERS-CoV, and SARS-CoV-2 produce an RDV-TP selectivity value less than 1 (~0.3). Effective inhibitory concentrations (EC_50_) measured in cell-based assays demonstrate potent antiviral activity of RDV across virus families. EC_50_ values for RDV or its parent nucleoside (GS-441524) are in the submicromolar range (Table 1). Here we demonstrate that HCV RdRp incorporates RDV-TP with a low selectivity value (0.93), indicating that both RDV-TP and ATP are used with similar efficiency. The published EC_50_ values between ~0.08 μM for RDV and ~4 μM for GS-441524 are also indicative of efficient antiviral activity although the prodrug has greater activity. A similar range of EC_50_ values is observed with RDV against the non-segmented negative-sense RNA viruses EBOV (0.003-0.021 μM), RSV (0.021 μM), and NiV (0.029-0.047 μM). The selectivity values are between 1.6 and 4, which are slightly higher when compared with our measurements for polymerases from the positive-sense RNA viruses. Conversely, RDV does not show significant antiviral activity against segmented RNA viruses CCHFV and FluB, and the EC_50_ value for LASV is relatively high (~4.5 μM for RDV). EC_50_ values have not been reported for GS-441524. In each of these cases, we also measured high selectivity values for RDV-TP incorporation, which are 20, 41, and 68, for LASV, CCHFV, and FluB, respectively. Thus, the combined results revealed a correlation between antiviral activity as measured in cell-based assays and efficient rates of incorporation of RDV-TP as measured in enzymatic assays. However, the incorporation of a nucleotide analogue does not necessarily translate into inhibition of RNA synthesis. The overarching remaining question is whether a uniform mechanism of action may help to explain the broad spectrum of antiviral activities associated with RDV.

### Inhibition of RNA synthesis catalyzed by HCV RdRp

Previous biochemical and structural studies with SARS-CoV-2 RdRp have shown that the incorporated RDV-MP causes delayed chain-termination due to a steric clash between 1’-cyano group and the hydroxyl group of the conserved Ser-861(26,27,29,31,32). We further demonstrated that a Ser-861-Gly mutant enzyme eliminated this blockage. A structural comparison of SARS-CoV-2 and HCV identified Gly-410 of the HCV RdRp as the residue equivalent to Ser-861 in SARS-CoV-2 RdRp (Fig. S1). Delayed chain-termination is therefore not expected with HCV RdRp. For SARS-CoV-2, the extent of inhibition at position 9, or “i+3” depends crucially on the concentration of the nucleotide substrate at position “i+4”. Low NTP concentrations favor inhibition, while high NTP concentrations override inhibition due to enhanced enzyme translocation. To monitor inhibition caused by RDV-TP, we utilized RNA templates with single sites of incorporation for RDV-TP and gradually increased the concentrations of the next NTP substrates (Fig. 1). Incorporation of RDV-TP by SARS-CoV-2 RdRp resulted in delayed-chain termination at position 9 (i+3) (Fig.1*B*). For HCV RdRp, subtle inhibition of RNA synthesis was observed at the point of incorporation (position 6, or i) and no inhibition at position 9, or “i+3” (Fig. 1*C*). Inhibition at position “i” is however easily overcome with low concentrations of the following NTP substrates. 50% read-through beyond the point of inhibition required a 10-fold greater nucleotide concentration for SARS-CoV-2 delayed chain-termination as compared to HCV (Fig. 1*D*).

**Figure 1.**
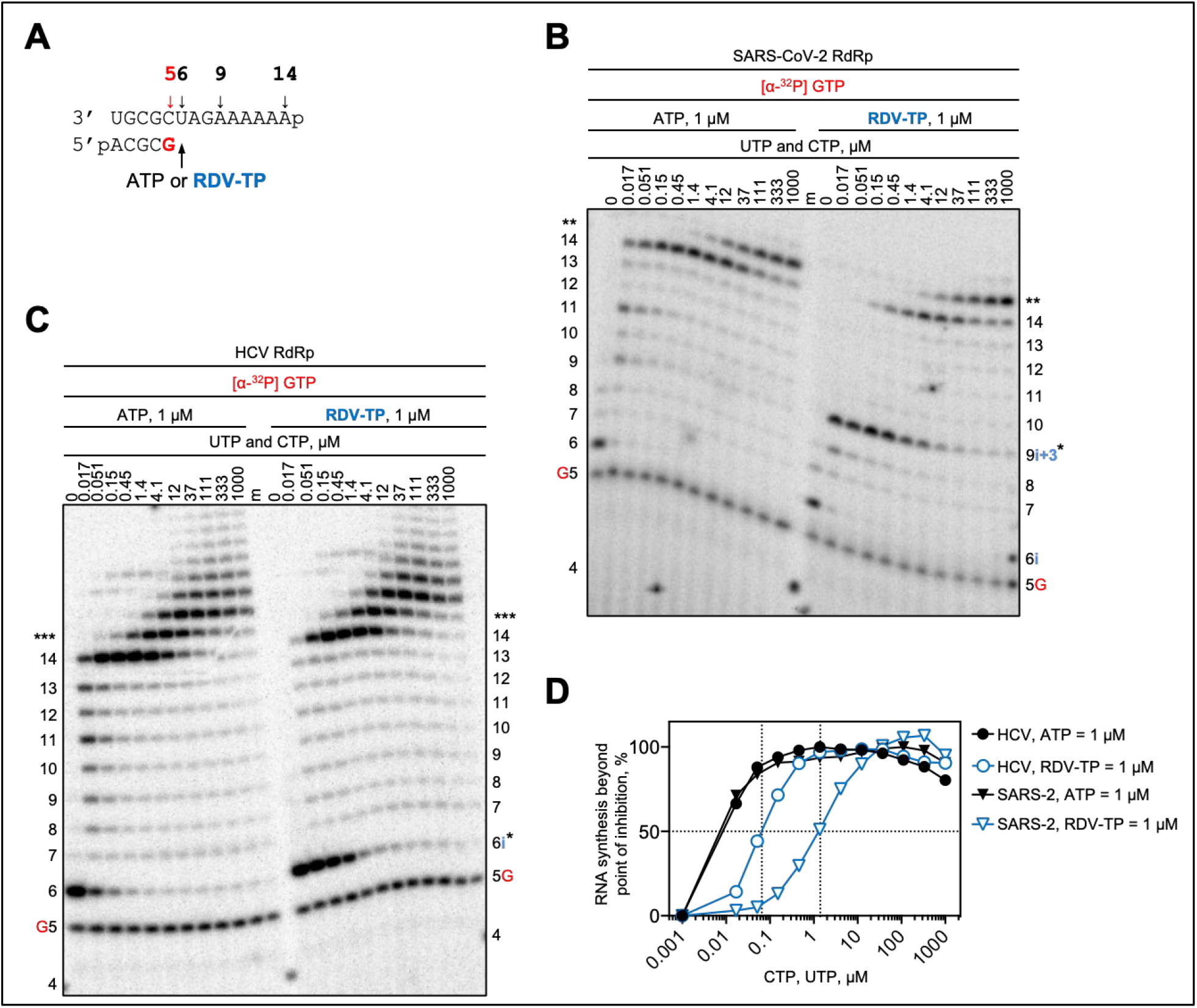
SARS-CoV-2 or HCV RdRp-catalyzed RNA synthesis and inhibition patterns following a single incorporation of RDV-MP as a function of nucleotide concentration. **(A)** RNA primer/template supporting a single incorporation event of AMP or RDV-MP at position 6. G indicates incorporation of [α-^32^P]-GTP at position 5. **(B)** Migration pattern of the products of RNA synthesis catalyzed by SARS-CoV-2 RdRp. A 5′-^32^P-labeled 4-nt primer (4) serves as a size marker. The asterisk indicates the point at which RNA synthesis is inhibited (position 9 or “i+3”). Two asterisks indicate slippage products that are likely sequence-dependent. **(C)** Reactions with HCV RdRp, inhibition of RNA synthesis occurs at the site of RDV-MP (position 6 or “i”). Three asterisks indicate increased slippage, likely sequence- and nucleotide-dependent. **(D)** Graphical representation of RNA synthesis beyond the point of RDV-TP induced inhibition for SARS-CoV-2 RdRp (“i+3”) and HCV RdRp (“i”).

To study whether repeated incorporations of RDV-TP could increase overall inhibition, we utilized a template with longer stretches of uridines (Fig. 2*A*). RNA synthesis by SARS-CoV-2 and HCV RdRp was monitored under competitive conditions in which we utilized a constant concentration of ATP and increasing concentrations of RDV-TP. RNA synthesis by SARS-CoV-2 RdRp was inhibited at ratios of RDV-TP over ATP as low as 0.1 and as high as 25 (Fig. 2*B*, *left*). An apparent rescue of RNA synthesis is seen at higher ratios of inhibitor over the natural nucleotide. This observation results from efficient RDV-TP incorporation by the SARS-CoV-2 RdRp that overrides inhibition; however, these high ratios of RDV-TP over ATP are unlikely relevant in a cellular environment. The possibility for multiple incorporations of RDV-TP by HCV RdRp did not show any significant RNA synthesis inhibition. (Fig. 2*B*, *right*). These results demonstrate that while SARS-CoV-2 and HCV RdRp both incorporate RDV-TP with high efficiency, significant inhibition of primer extension reactions is solely seen with SARS-CoV-2 RdRp. While RDV-mediated delayed chain-termination may therefore contribute to the observed antiviral effects in coronaviruses, the antiviral effect of RDV against HCV is likely based on a different mechanism.

**Figure 2.**
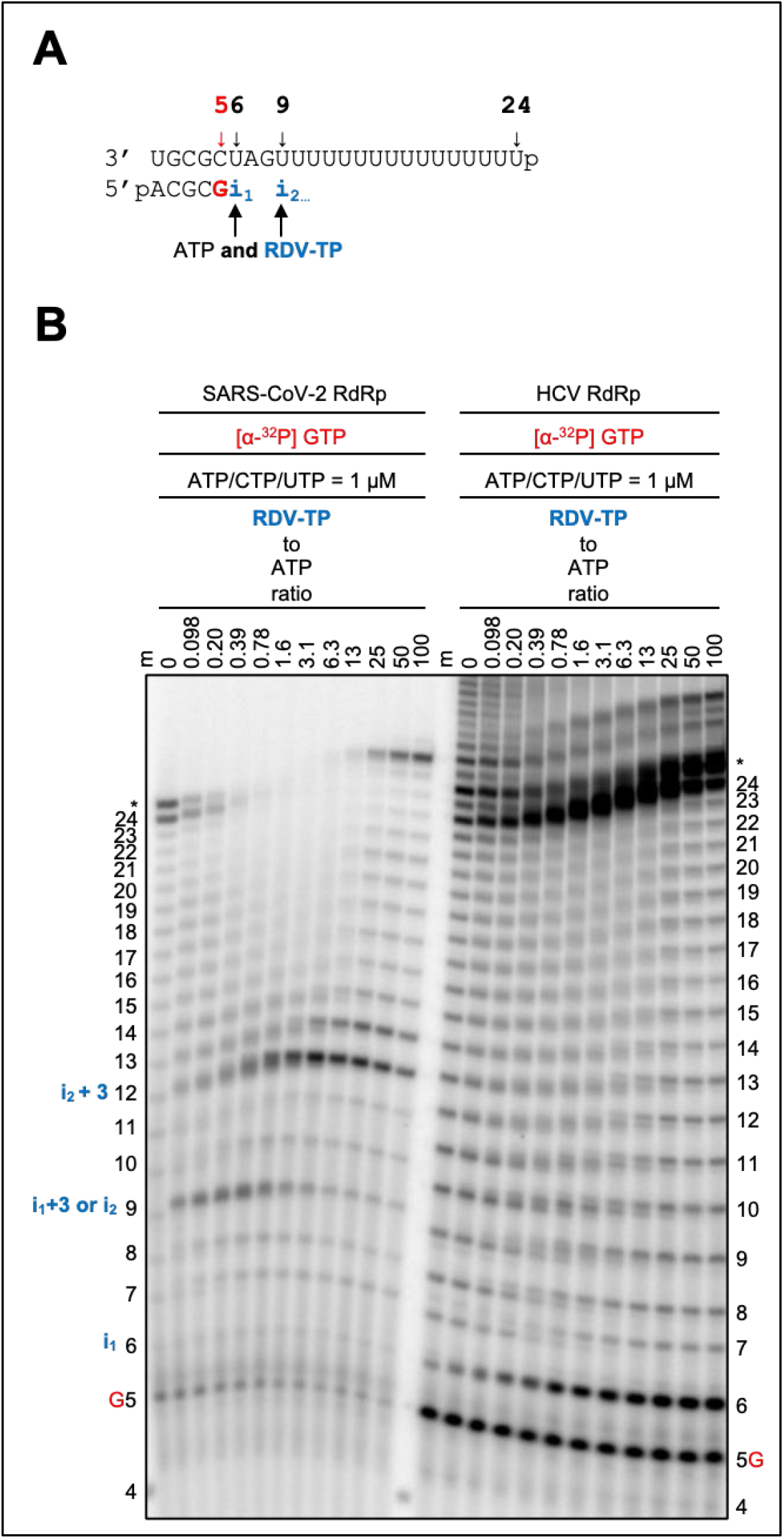
SARS-CoV-2 or HCV RdRp-catalyzed RNA synthesis following multiple incorporations of RDV-MP as a function of RDV-TP concentration in the presence of a constant NTP concentration. **(A)** RNA primer/template supporting multiple incorporation events of ATP or RDV-TP. **(B)** Migration pattern of the products of RNA synthesis catalyzed by SARS-CoV-2 RdRp (left), First and second incorporation of RDV-TP occurs at position 6 (“i“) and 9 (“i_2_”), respectively, with inhibition appearing at position 9 (“i+3”) and 12 (“i_2_+3”). Migration pattern of HCV RdRp (right), minor product accumulation occurs at position 6, but inhibition of RNA synthesis resulting in full template length product is not evident.

### Template-dependent inhibition of HCV RNA synthesis

High rates of incorporation of RDV-TP along with mechanisms that overcome inhibition provide conditions that allow synthesis of full-length RNA copies. These copies are modified with multiple RDV-MP residues that could affect RNA synthesis when used as templates. For SARS-CoV-2 RdRp, the RDV-MP in the template causes inhibition of the incorporation of complementary UTP and the following NTP. A similar mechanism is also considered for HCV RdRp (Fig. 3). We used RNA templates with a single incorporated RDV-MP at position 11. Reactions were conducted with increasing concentrations of UTP to determine a threshold of inhibition. Both SARS-CoV-2 and HCV RdRp show inhibition at position 10 prior to the site of UTP incorporation. Increasing UTP concentration diminishes inhibition in both cases. HCV RdRp also requires a lower concentration of UTP to overcome this obstacle. While inhibition of SARS-CoV-2 RdRp is also seen at the adjacent position, inhibition of HCV is confined to position 10. As a result of these two effects, overall inhibition of RNA synthesis is more pronounced for SARS-CoV-2 RdRp. In both cases, template-dependent inhibition is more pronounced than the inhibitory effects in primer extension reactions.

**Figure 3.**
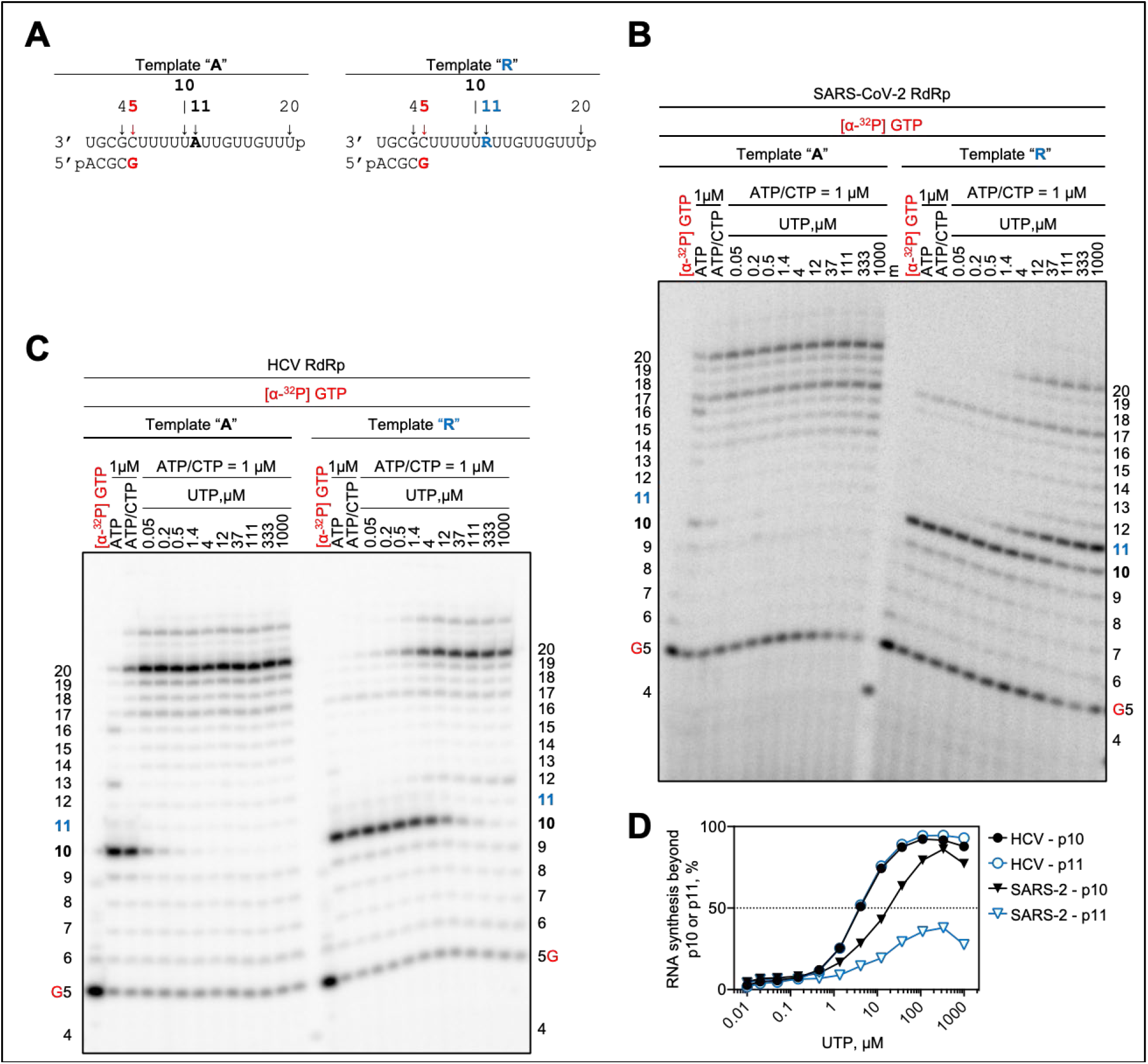
Template-dependent inhibition of SARS-CoV-2 and HCV RdRp by an embedded RDV-MP **(A)** RNA primer/template with an embedded AMP (Template “A”, *left*) and RDV-MP (Template “R”, *right*) at position 11. **(B)** Migration pattern of products of RNA synthesis catalyzed by SARS-CoV-2 RdRp, incorporation of ATP at position 12 is inhibited **(C)** Reactions with HCV RdRp, incorporation of ATP at position 12 is not inhibited. **(D)** Graphical representation of RNA synthesis beyond position 10 and 11 (p10 and p11, respectively) on template “R”.

### Embedded RDV-MP demonstrates a unified mechanism of inhibition

RDV-dependent delayed chain termination had been previously observed in the context of NiV RdRp and RDV-TP, albeit following multiple subsequent RDV-TP incorporations(11). NiV RdRp RNA synthesis was here monitored following a single RDV-TP or ATP incorporation in conjunction with increasing NTP concentrations (Fig. 4*A*). However, we were unable to detect significant inhibitory effects under these conditions. Similar intermediate products were consistent beyond AMP and RDV-MP incorporation and can likely be attributed to low processivity. Also, no significant inhibition immediately following RDV-TP incorporation suggests that the newly synthesized template strand contains RDV-MP residues embedded throughout. Employing the same approach as above, RNA synthesis inhibition was again seen opposite RDV-MP at position 11 (Fig. 4*B*). Incorporation opposite templated RDV-MP and AMP, respectively, was evaluated with increasing concentrations UTP. Templated AMP allowed RNA synthesis to proceed to the full-length RNA at a UTP concentration as low as 4 μM. In contrast, a UTP concentration up to 1000 μM was not sufficient to support incorporation opposite RDV-MP, thus halting RNA synthesis at position 10 (Fig. 4*B*). Previous work with RSV and EBOV RdRp revealed difficulties in identifying a template that enables the investigation of a single RDV-TP incorporation and generation of full template length product(16,25). However, multiple incorporations of RDV-TP can cause delayed chain-termination with RSV and EBOV RdRp. Here we also examined RNA synthesis opposite the embedded RDV-MP at position 11 of the template (Fig. S2). RSV RdRp catalyzed RNA synthesis could not proceed beyond the embedded RDV-MP, with most of the product accumulating at position 10, even at the highest NTP concentrations (Fig. S2, *A*). The diminished processivity of EBOV RdRp makes it difficult to accurately determine RNA synthesis and its inhibition beyond 10 nucleotides (Fig. S2, *B*).

**Figure 4.**
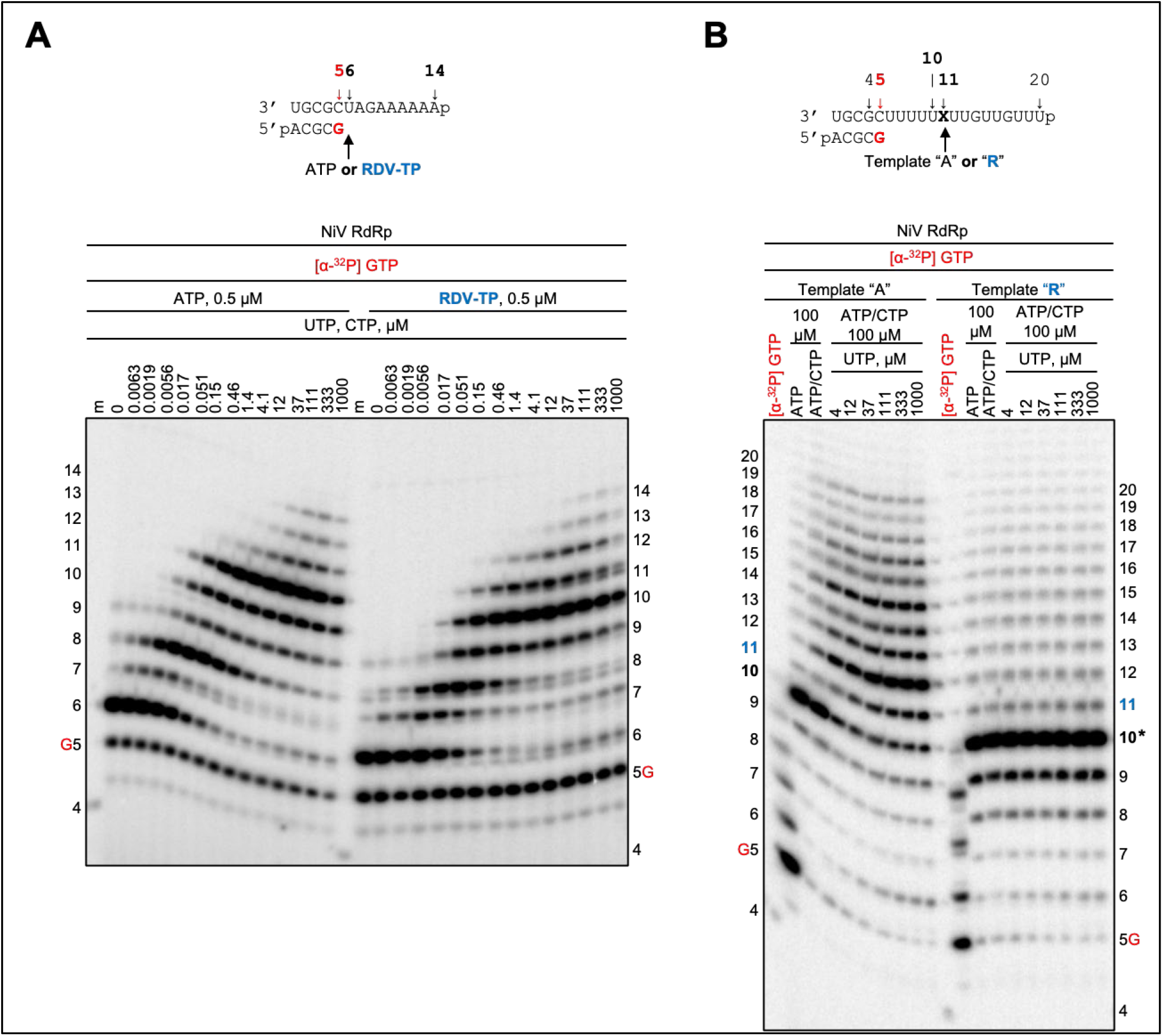
RNA synthesis patterns following AMP and RDV-MP incorporation and template-dependent inhibition of NiV RdRp. (**A**) RNA primer/template as indicated in Fig. 1 and migration pattern of products of RNA following the incorporation of AMP or RDV-MP. No inhibition is evident. **(B)** RNA primer/template as indicated in Fig. 3 and migration pattern of RNA synthesis opposite AMP (*left*) or RDV-MP (*right*). The asterisk at position 10 indicates the point of inhibition as a result of the embedded RDV-MP at position 11.

Finally, we studied the inhibitory effects of RDV-TP or RDV-MP against RdRp enzymes from segmented negative-sense RNA viruses. We have previously demonstrated that RDV-TP is also a weak substrate for LASV RdRp and its incorporation does not cause significant inhibition (26). Here we demonstrate that the embedded RDV-MP in the template causes a complete stop of RNA synthesis (Fig. S2*C*). For FluB RdRp, the presence of RDV-TP also does not mediate significant inhibition of RNA synthesis (Fig. 5*A*). Minor differences in the degree of full-length product formed can likely be attributed to the inefficient incorporation of RDV-TP as compared to ATP. Templates containing AMP and RDV-MP, respectively, demonstrate again that the embedded RDV-MP causes RNA synthesis arrest at position 10 (Fig. 5*B*). A very similar pattern is seen with CCHFV RdRp (Fig. 6*A*/*B*): no significant inhibition by RDV-TP in primer extensions, and strong inhibition or even termination when RDV-MP is embedded in the template. However, inhibition translates to antiviral effects only if the rate of incorporation of RDV-TP is sufficiently high.

**Figure 5.**
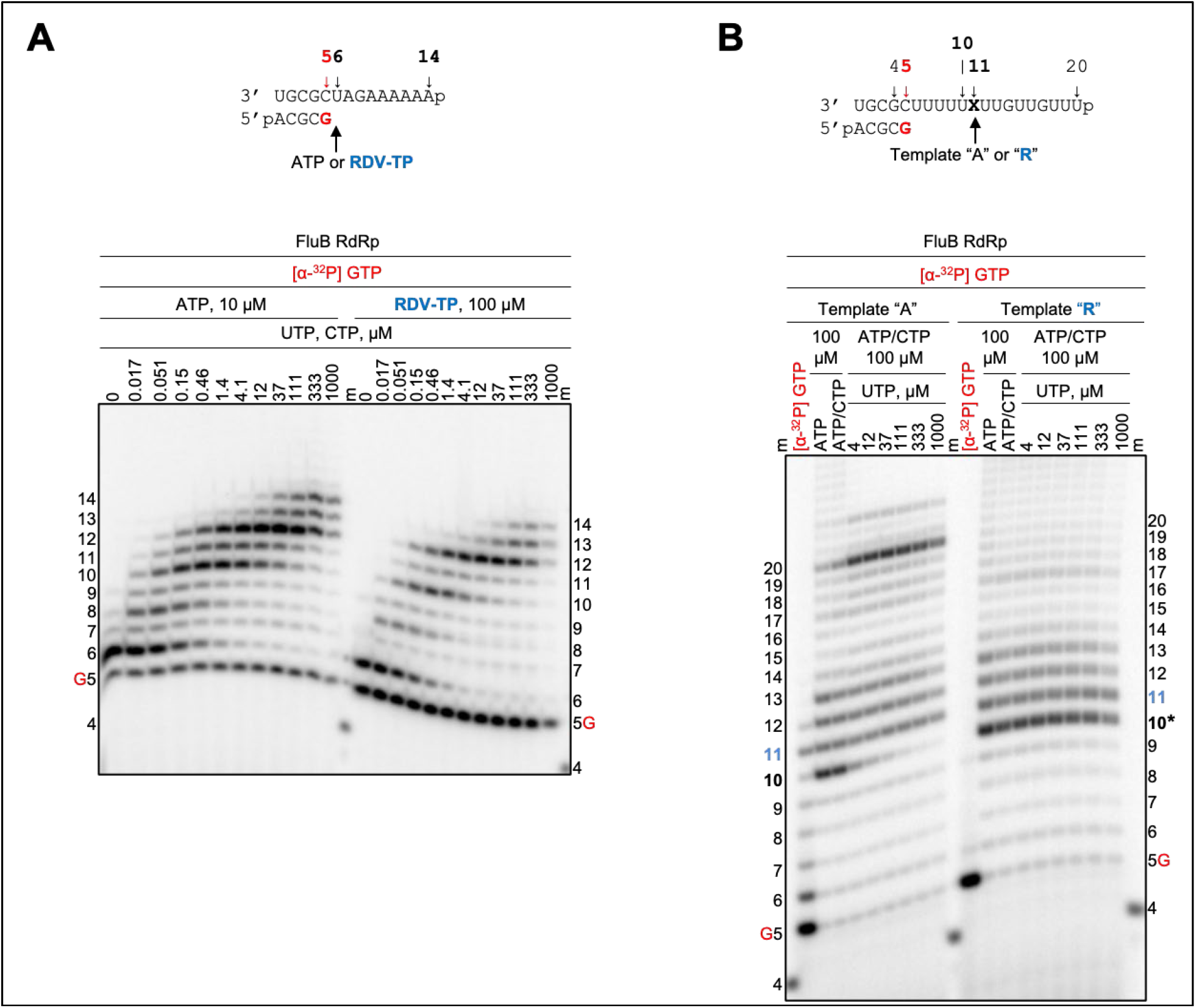
RNA synthesis patterns following AMP and RDV-MP incorporation and template-dependent inhibition of FluB RdRp. (**A**) RNA primer/template as indicated in Fig. 1 and migration pattern of RNA products following the incorporation of AMP or RDV-MP. No inhibition is evident. **(B)** RNA primer/template as indicated in Fig. 3 and migration pattern of RNA synthesis opposite AMP (*left*) or RDV-MP (*right*). The asterisk at position 10 indicates the point of inhibition as a result of the embedded RDV-MP at position 11.

**Figure 6.**
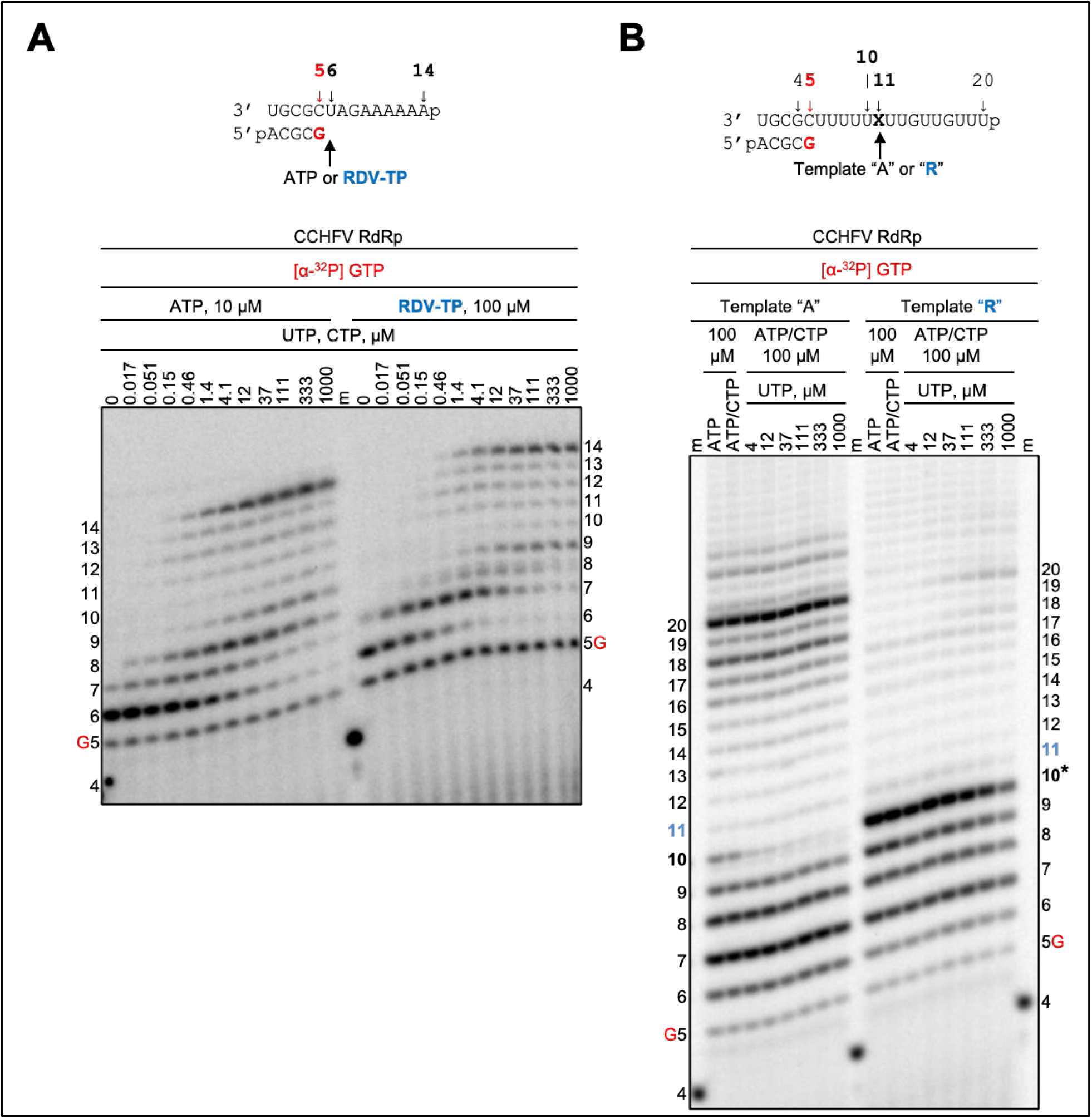
RNA synthesis patterns following AMP and RDV-MP incorporation and template-dependent inhibition of CCHFV RdRp. (**A**) RNA primer/template as indicated in Fig. 1 and migration pattern of products of RNA following the incorporation of AMP or RDV-MP. No inhibition is evident. **(B)** RNA primer/template as indicated in Fig. 3 and migration pattern of RNA synthesis opposite AMP (*left*) or RDV-MP (*right*). The asterisk at position 10 indicates the point of inhibition as a result of the embedded RDV-MP at position 11.

## Discussion

We compared the inhibitory effects of RDV-TP against a panel of viral RNA polymerases that represent diverse families of RNA viruses. The goal was to identify correlations between antiviral effects as previously measured in cell-based assays and the mechanism of inhibition. Members of the positive sense *Coronoviridae* (SARS-CoV-2) and *Flaviviridae* (HCV) families, as well as the non-segmented negative sense *Pneumoviridae* (RSV), *Filoviridae* (EBOV) and *Paramyxoviridae* (NiV) families are sensitive to RDV treatment, while members of the segmented negative sense *Arenaviridae* (LASV), *Orthomyxoviridae* (FluB), and *Nairoviridae* (CCHFV) families are not sensitive to the drug (8,13,19,20). We expressed the corresponding RdRp enzymes or enzyme complexes and studied: (i) selective incorporation of the nucleotide substrate RDV-TP, (ii) the effect of the incorporated RDV-MP on primer extension, and (iii) the effect of the template-embedded RDV-MP on UTP incorporation.

Efficient selective incorporation of RDV-TP, calculated as ratio of efficiency of incorporation of ATP over RDV-TP, is seen with SARS-CoV-2 (0.3), HCV (0.9), EBOV (4.0), NiV (1.6) and RSV enzymes (2.7). In contrast, much higher selectivity values for LASV (20), CCHFV (41), and FluB (68) enzymes suggest poor RDV-TP substrate usage. The ability of a nucleotide analogue inhibitor to be incorporated by viral RdRp enzymes is largely dependent on variations in the residues that define the nucleotide binding site(33). The overall structure of the active site is well conserved across a wide array of viruses and is commonly defined by a set of motifs(34). Of these, motifs A, B, and C form important interactions with the incoming NTP.

From available ternary structures of SARS-CoV-2, HCV and FluB RdRp, primer/template and incoming NTP, we developed models of how RDV-TP binds in its pre-incorporated state as compared to ATP (Fig. 7). Consistent with our biochemical findings, SARS-CoV-2 and HCV RdRp enzymes show similar active sites for favorable binding RDV-TP with virtually no distortion in position or alteration in how the ribose is recognized as compared to ATP. Moreover, the 1’-pocket is polar and therefore conducive to accommodating the 1’-cyano group of RDV-TP. In contrast, for FluB RdRp, polar residues at the active site appear to be too far to interact directly with the ribose portion of the NTP, as in SARS-CoV-2 and HCV. Moreover, a WaterMap analysis predicted the presence of a water molecule in the nucleotide binding site (35,36). The location of this water molecule overlaps with the location of the 1’-cyano group and would be necessarily displaced. A sequence alignment of FluB, LASV and CCHFV enzymes suggests similarities, which may explain the poor incorporation of RDV-TP by these polymerases. Structures of EBOV, NiV, CCHFV RdRp enzymes are not available, and structure of RSV and LASV RdRp lack the RNA substrate, which is a limitation of our modeling approach.

**Figure 7.**
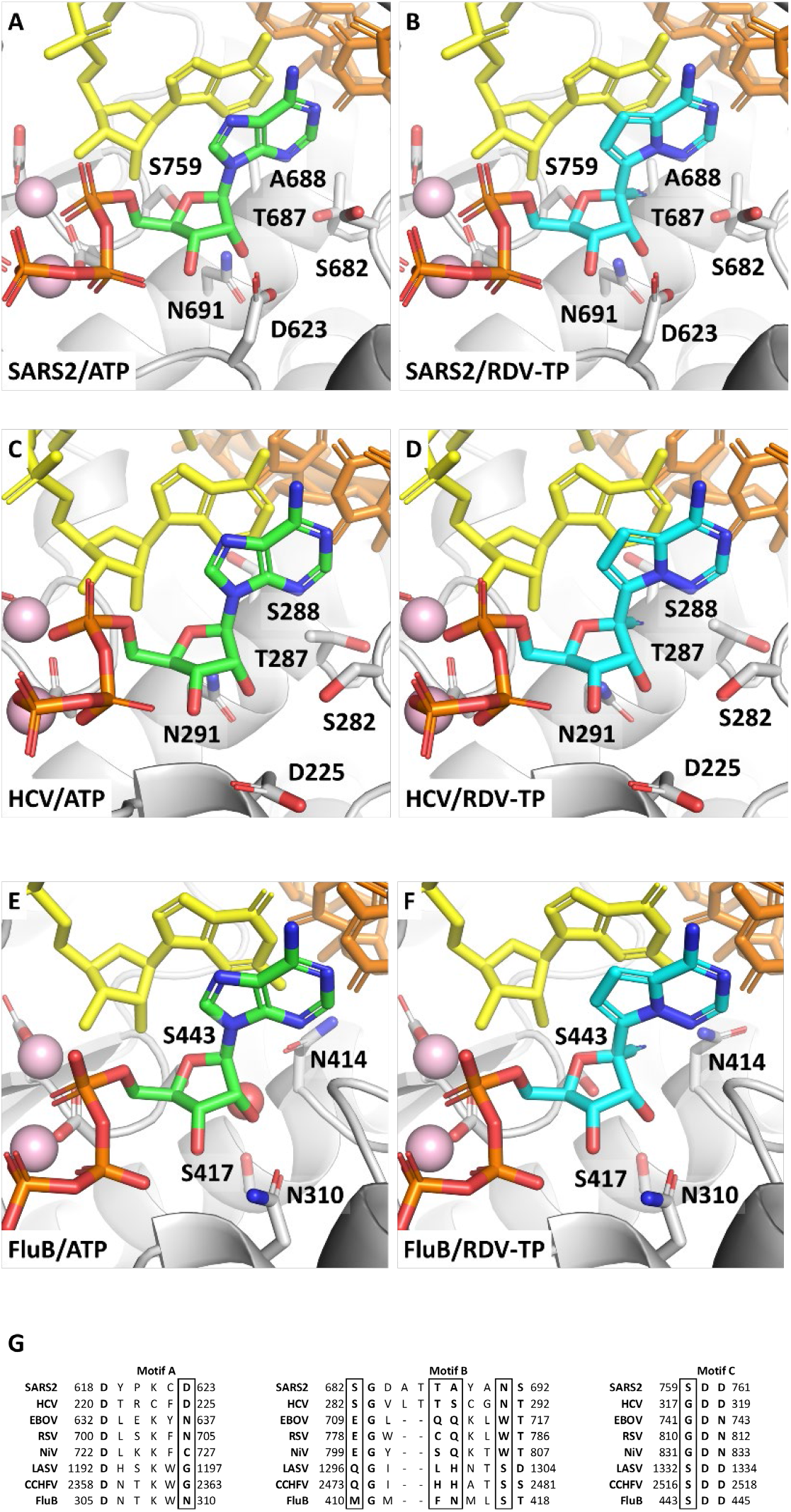
Models of ATP and RDV-TP in their pre-incorporation states for SARS-CoV-2 **(A,B)**, HCV **(C,D)** and FluB **(E,F)** RdRps. RDV-TP is seen to be a good substrate for SARS-CoV-2 and HCV, showing little difference in binding position relative to ATP. However, RDV-TP displaces a water molecule critical to recognition of the ribose 2’OH in FluB, compromising its binding affinity. **(G)** A comparison of the key motifs that make up the polymerase active site suggests LASV and CCHFV may recognize the NTP in a manner more similar to FluB, while EBOV and RSV likely recognize the NTP by a different set of interactions.

Clear evidence for delayed chain-termination following a single incorporation of RDV-TP has so far only been demonstrated for SARS-CoV, MERS-CoV, and SARS-CoV-2 RdRp complexes. In these cases, a conserved serine clashes with the 1’-cyano group of the incorporated RDV-MP at position “i+3”(26,27,29,31,32). However, NTP concentrations > 10 μM are often sufficient to overcome the inhibition(26,27,29,31). The clash between the 1’-cyano group of RDV-MP and S861 in SARS-CoV-2 occurs in a region of the RdRp away from the active site. Indeed, tested polymerases of other viruses did not show any significant inhibition in primer extension reactions, although previous studies have shown that multiple consecutive incorporations of the inhibitor can also lead to delayed chain-termination at positions “i+3” and “i+5” with EBOV, RSV, and NiV RdRp complexes(11,16). In this work, we have also shown a subtle inhibitory effect at position “i” with HCV RdRp. Overall, this analysis suggests that the inhibitory effect in primer extension reactions is heterogeneous and generally weak. In contrast, the template-dependent inhibition of RNA synthesis seems to provide a unifying mechanism that shows strong inhibition of UTP incorporation opposite RDV-MP.

The template nucleotide that base-pairs with the incoming nucleotide substrate is positioned through conserved interactions with residues in motif F (Fig 8). In particular, the template base interacts with a bulky hydrophobic residue, while the ribose interacts with a second residue of varying character. These residues are separated by one additional residue which is turned away from the template. In the case of SARS-CoV-2, the base moiety contacts V557 while the ribose interacts with G559. For HCV, similar interactions are seen with I160 and Y162, while for FluB, the equivalent interactions are with I241 and T243. A sequence alignment suggests that this sub-motif is conserved across all enzymes included in this study. In each case, the template RDV-MP 1’-cyano would be positioned between these two motif F residues, having a potential clash with the backbone of the middle residue. In SARS-CoV-2, the 1’-cyano is positioned too close to the carbonyl of A558 (29). Similarly, the 1’-cyano appears too close to the carbonyl of A242 in FluB. In addition, the hydrophobic nature of this area is likewise not conducive to placement of the polar cyano group and may also drive a shift in the position of template RDV.

**Figure 8.**
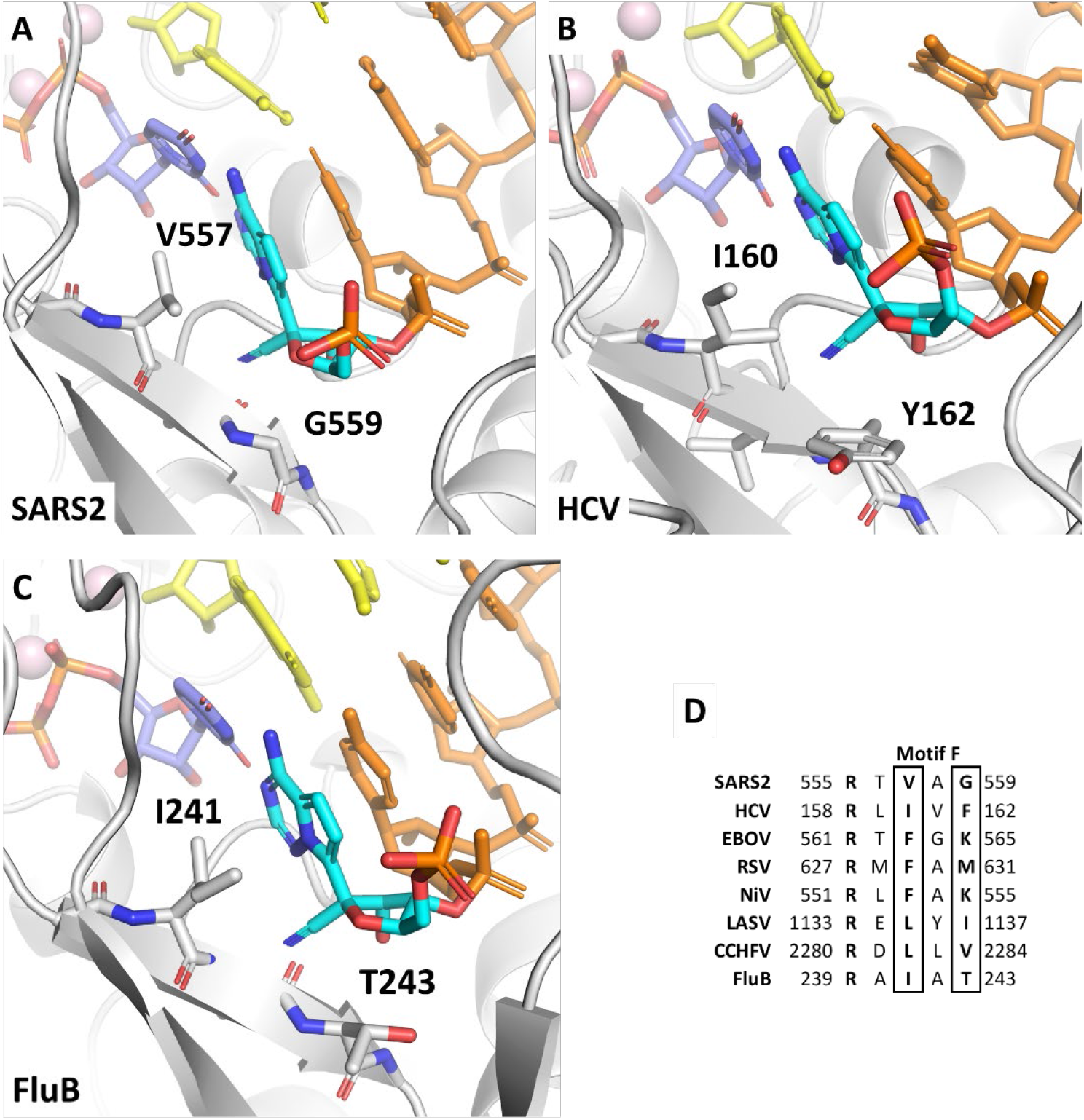
Template-dependent inhibition by RDV is shown here for **(A)** SARS-CoV-2, **(B)** HCV and **(C)** FluB RdRps. The 1’-CN of template incorporated RDV-MP is positioned between two typically hydrophobic residues, pushing against the backbone of the intermediate residue. **(D)** As the overall structure of Motif F is widely conserved, if not the exact sequence, this represents a uniform mechanism of inhibition across a diverse set of polymerases.

Taken together, the combined results of this biochemical study and previously reported cell-based data suggest a strong correlation between the antiviral effect of RDV, or its nucleoside parent, and effective use of RDV-TP as substrate. The high selectivity of RDV-TP over ATP is a prerequisite for any downstream inhibitory effect. The patterns of chain-termination or delayed chain-termination can differ substantially depending on the nature of the polymerase. However, inhibition is usually inefficient at higher NTP concentrations that facilitate polymerase translocation and continuation of RNA synthesis. For coronaviruses, weak inhibition during synthesis of the first RNA strand might even be desired, so as to evade the intrinsic proofreading activity associated with the replication complex. A uniform mechanism of action is provided by the template-dependent inhibition of RNA synthesis opposite the embedded RDV-MP. Inhibition of incorporation of an UTP base-pairing with RDV-MP is likely based on steric conflicts between the 1’-cyano group of RDV-MP and conserved residues of motif F. This inhibitory effect is seen with each of the diverse RdRp enzymes used in this study, including polymerases from viruses that are not sensitive to RDV. In these cases, the selective use of RDV-TP as substrate was poor. Thus, future drug development efforts that aim at a broader spectrum of antiviral activities should focus on improving rates of incorporation while still capitalizing on the inhibitory effects of a bulky 1’-modification.

## Materials and Methods

### Nucleic acids and chemicals

RNA primers and templates used in this study were 5′-phosphorylated and purchased from Dharmacon (Lafayette, CO, USA). RDV-TP was provided by Gilead Sciences (Foster City, CA, USA). NTPs were purchased from GE Healthcare. [α-^32^P]GTP was purchased from PerkinElmer.

### Protein expression and purification

Expression and purification of RNA-dependent RNA polymerases (RdRp) used in this study have been described (11,25,26,29,30,37). We also utilized the baculovirus expression system for HCV nsp5b RdRp and NiV RdRp P/L complex. The pFastBac-1 (Invitrogen, Burlington, ON, Canada) plasmid with the codon-optimized synthetic DNA sequences (GenScript, Piscataway, NJ, USA) coding for HCV nsp5b RdRp (CAB46677.1 polyprotein, residues 2420 to 3010) minus the 21 C-terminus residues was used as a starting material for protein expression in insect cells (Sf9, Invitrogen, Burlington, ON, Canada). We employed the MultiBac (Geneva Biotech, Indianapolis, IN, USA) system for protein expression in insect cells (Sf9, Invitrogen, Burlington, ON, Canada) according to published protocols (38,39). HCV nsp5b RdRp was purified using Ni-NTA affinity chromatography based on the C-terminal eight-histidine tag according to the manufacturer’s specifications (Thermo Scientific, Rockford, IL, USA). NiV RdRp P/L complex coding sequence (AEZ01385 and AEZ01390, P and L, respectively) was expressed in insect cells as a polyprotein in frame with N-terminal Tobacco etch virus (TEV) protease. This approach was originally reported for the expression of influenza virus RdRp trimeric complex(40,41). NiV P and L proteins were cleaved post-translationally from the polyprotein at the engineered TEV sites. NiV RdRp P/L complex was purified using Ni-NTA affinity chromatography based on the P protein N-terminal eight-histidine tag according to the manufacturer’s specifications (Thermo Scientific, Rockford, IL, USA). The protein identities of the purified HCV nsp5b RdRp and NiV RdRp P/L complex were confirmed by mass spectrometry (MS) analysis (Alberta Proteomics and Mass Spectrometry, Edmonton, AB, Canada).

### Evaluation of nucleotide incorporation and the effect of primer- or template-embedded remdesivir on viral RNA synthesis

The following synthetic 5’-monophosphorylated RNA templates were used in this study (the portion of the template which is complementary to the 4-nt primer is underlined): 3’<u>UGCG</u>CUAGAAAAAAp for measurements of the RDV-TP selectivity values and determination of the patterns of inhibition of RNA synthesis subsequent to an incorporated RDV at single position; 3’<u>UGCG</u>CUAGUUUUUUUUUUUUUUUUp for ATP/RDV-TP competition experiments; 3’<u>UGCG</u>CUUUUURUUGUUGUUUp for evaluation of nucleotide incorporation opposite template-embedded remdesivir (R, template “R”) in comparison to templated adenosine at the equivalent position (template “A”). Both RNA templates were produced in the lab as previously described by us (29). RNA synthesis assays of viral RdRp, data acquisition and quantification were done as previously reported by us (16,25,26,29,30). Briefly, concentrations of various viral RdRp proteins and protein complexes were selected such that incorporation of [α-^32^P]-GTP was linear during single nucleotide incorporation studies or viral enzyme concentrations were optimized to promote full template-length RNA synthesis involving primer- or template-embedded remdesivir. Standard reaction mixture for RNA synthesis assays (final concentrations after mixing) included optimal concentrations of purified viral RdRp, Tris-HCl (pH 8, 25 mM), RNA primer (200 μM), RNA template (2 μM, except for FluB RdRp reactions where optimal RNA template concentration was 0.5 μM), [α-^32^P]-GTP (0.1 μM), various concentrations and combinations (as indicated) of NTP and RDV-TP. Partial (minus MgCl2) reaction mixtures (10 μL) were prepared on ice and incubated for 5-10 minutes at 30 °C followed by the addition of 5 μL of MgCl2 (5 mM) to initiate nucleotide incorporation. Reactions were stopped after 30 min by the addition of 15 μL of formamide/EDTA (50 mM) mixture and incubated at 95 °C for 10 min. Reaction products were resolved through 20 % PAGE, and the [α-^32^P]-generated signal was stored and scanned from phosphorimager screens. Data were analyzed using GraphPad Prism 7.0 (GraphPad Software, Inc., San Diego, CA, USA).

### Molecular Modelling

To date, a well resolved pre-incorporation state for SARS-CoV-2, in which the NTP is positioned in the active site with suitable primer and template RNA, has not been determined by cryo-EM. A model of the pre-incorporation state of RDV-TP in SARS-CoV-2 has previously been described (26,29). Based on more recent cryo-EM structures, including PDB:6XEZ (1), which captures the SARS-CoV-2 polymerase complex (nsp12/nsp7/(nsp8)2/(nsp13)2) with primer and template RNA bound, the model has since been updated using similar approaches to those outlined previously. Models for HCV were based on the x-ray structures PDB:4WTA and PDB:4WTD (42), which capture pre-incorporation states of UDP and ADP, respectively. Models for FluB were based on the cryo-EM structures PDB:6QCV and PDB:6QCT (43), which capture the pre-incorporation initiation state and the elongation state, respectively. All optimization was performed with Macromodel and Prime in the Schrödinger Suite (44,45). The identification of a water molecule in the active site of FluB, which establishes a hydrogen bond network between the protein and the NTP, was done with WaterMap, also part of the Schödinger Suite (35,36).

## Data availability

All data are included within this article

## Supporting information

This article contains Supporting information

## Acknowledgements

The authors would like to thank Emma Woolner and Dana Kocincova for excellent technical assistance and Dr. Jack Moore at the Alberta Proteomics and Mass Spectrometry facility for mass spectrometry analysis.

## Funding and additional information

This study was supported by grants to MG from the Canadian Institutes of Health Research (CIHR, grant number 170343), Gilead Sciences and from the Alberta Ministry of Economic Development, Trade and Tourism by the Major Innovation Fund Program for the AMR – One Health Consortium.

## Abbreviations and nomenclature

RDV: remdesivir
COVID-19: coronavirus disease 2019
SARS-CoV-2: severe acute respiratory syndrome coronavirus 2
HCV: hepatitis C virus
NiV: Nipah virus
Flu: influenza
FluA: influenza A
FluB: influenza B
CCHFV: Crimean-Congo hemorrhagic fever virus
RdRp: RNA-dependent RNA polymerases
TP: triphosphate
MP: monophosphate
WHO: World Health Organization
EVD: Ebola virus disease
MERS-CoV: Middle Eastern respiratory syndrome
EBOV: Zaire ebolavirus
RSV: respiratory syncytial virus
LASV: Lassa virus
EC_50_: effective inhibitory concentration

