## Supplemental Information for "Efficient incorporation and template-dependent polymerase inhibition are major determinants for the broad-spectrum antiviral activity of remdesivir"

**Supplemental Figure 1.** Structure-based alignment of select CoV and HCV RdRps.
**Supplemental Figure 2.** Template-dependent inhibition of RSV, EBOV, and LASV RdRps by an embedded RDV-MP.

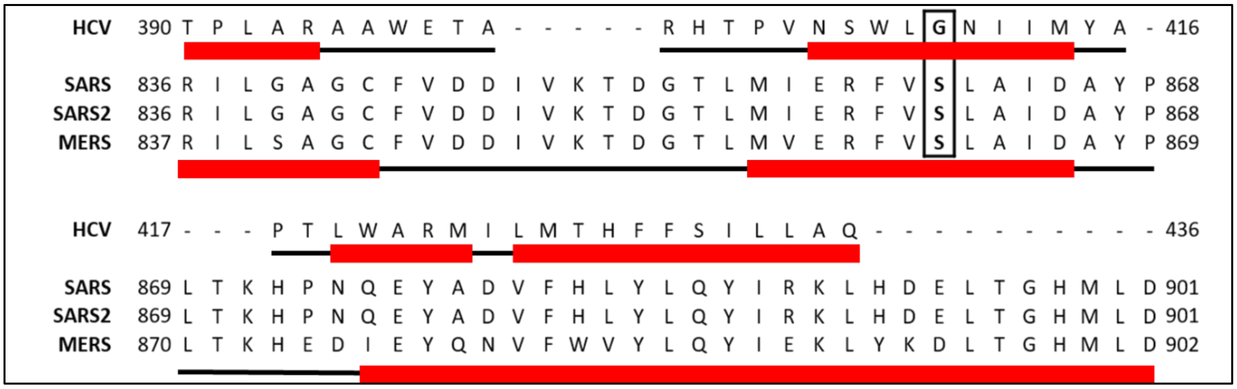


**Supplemental Figure 1.** Structure-based alignment of select CoV and HCV RdRps. Sequence alignment based on a 3D structural overlay of HCV ns5b (PDB:4WTA) and SARS-CoV-2 nsp12 (PDB:6XEZ). Secondary structure is shown below the HCV sequence and below the three coronavirus sequences. Helix secondary structure elements are indicated by the red bars, while loops are indicated by solid black lines. While the three coronaviruses have high homology, the HCV sequence has fairly low homology. Despite this, it shares a common secondary structure in the thumb sub-domain such that Ser-861 in SARS-CoV-2 is seen to correspond to Gly-410 in HCV.

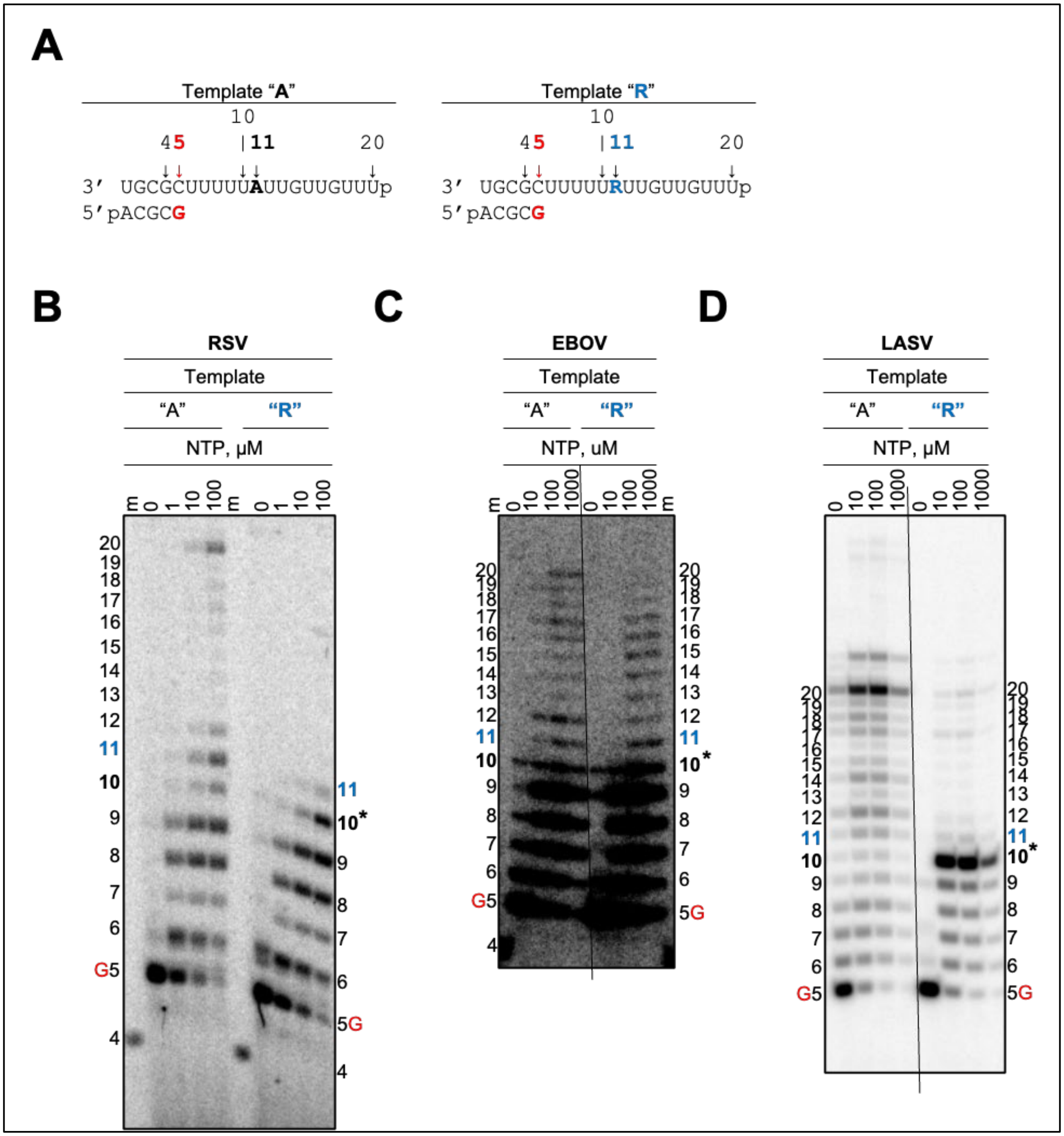


**Supplemental Figure 2.** Template-dependent inhibition of RSV, EBOV, and LASV RdRps by an embedded RDV-MP. **(A)** RNA primer/template as indicated in Fig. 3. **(B)** Migration pattern of RSV RdRp-catalyzed RNA synthesis supplemented with increasing NTP concentrations opposite AMP (*left*) or RDV-MP (*right*). **(C)** Migration pattern of EBOV RdRp-catalyzed RNA synthesis in which the conditions are the same as described in **B**. **(D)** Migration pattern of RSV RdRp-catalyzed RNA synthesis in which the conditions are the same as described in **B** and **C**. The asterisk at position 10 indicates point of inhibition as a result of the embedded RDV-MP at position 11.
